# Integrating Artificial Intelligence and Bioinformatics Methods to Identify Disruptive STAT1 Variants Impacting Protein Stability and Function

**DOI:** 10.1101/2024.10.04.616446

**Authors:** Ebtihal Kamal, Lamis A. Kaddam, Mehad Ahmed, Abdulaziz Alabdulkarim

**Affiliations:** Department of basic medical sciences,Faculty of medicine, Prince Sattam Bin Abdulaziz university,KSA; Department of Physiology, Faculty of Medicine King Abdul-Aziz University Rabigh, KSA; Plastic Surgery, Department of Surgery, College of Medicine, Prince Sattam Bin Abdulaziz University, Al Kharj, Saudi Arabia

**Author notes:** **Corresponding authors** Correspondence to Ebtihal Kamal.

**Keywords:** STAT1 gene, signalling pathway, single nucleotide polymorphisms, bioinformatics, artificial intelligence

## Abstract

The Signal Transducer and Activator of Transcription *1 (STAT1)* gene is an essential component of the JAK-STAT signalling pathway. This pathway has a pivotal role in regulating different cellular processes, including immune responses, cell growth, and apoptosis. Mutations in the *STAT1* gene contribute to various body pathologies [OMIM #600555], including immune system dysfunction.

In the current study, we used eighteen online computational approaches. Six pathogenic variants (R602W, I648T, V642D, L600P, I578N, and W504C) were predicted to significantly disrupt protein stability and function. These findings highlight the potential of approaches to pinpoint pathogenic single nucleotide polymorphisms, providing a time and cost effective alternative to experimental approaches. Up to the best of our knowledge, this is the original inclusive study, where we analyze *STAT1* gene variants using both bioinformatics and artificial intelligence based model tools.

## Introduction

The Signal Transducer and Activator of Transcription 1 (*STAT1*) gene is an essential mediator of the JAK-STAT signalling pathway in response to interferons ^[1–4]^. It plays a crucial role in the biological immune response against intracellular mycobacterial infections as well as viral infections ^[1,5,6]^. The *STAT1*gene in humans is composed of seven domains located on chromosome 2q32.2 and consists of 25 exons ^[1,7,8]^. Genetic variants within the *STAT1* gene lead to loss-of-function (LOF) and gain-of-function (GOF) phenotypes with a wide range of clinical presentations, including autoimmunity, life-threatening mycobacterial, severe viral and bacterial infections^[9–11]^[OMIM #600555].

GOF mutation is associated with chronic mucocutaneous candidiasis ^[12,13]^, while patients with LOF mutations display an increased susceptibility to intracellular bacteria, including a Mendelian susceptibility to mycobacterial disease (MSMD)^[2,14]^. Despite these mutations being well investigated, the role of single-nucleotide polymorphisms (SNPs) in STAT1-associated disorders is inadequately elucidated. (SNPs) constitute a common form of genetic variation in humans ^[15]^. The nonsynonymous SNPs (nsSNPs) cause alteration in the amino acid residues as a result of variation in the sequence of DNA at a single position of a nucleotide (A, T, C, or G) which contributes to the functional diversity of the related proteins ^[16,17]^.

Although bioinformatics tools have played a role in the prediction of damaging SNPs and their relationship with diseases ^[18]^. The influence of STAT1 nsSNPs on protein function has not been thoroughly investigated, despite their potential importance, this indicates a substantial scientific gap. Nonetheless, no published research has systematically examined STAT1 SNPs by bioinformatics approaches. In this study, we will perform a comprehensive STAT1-SNPs analysis using bioinformatics prediction tools combined with artificial intelligence models to identify the pathogenic and deleterious SNPs, providing novel insights into their involvement in immune dysregulation and establishing a foundation for subsequent functional and clinical research.

## Results

### Distribution of STAT1 gene SNP datasets

The SNPs of the human STAT1 gene were collected from the NCBI (https://www.ncbi.nlm.nih.gov). The total number of SNPs was 10989; there were 480 SNPs located in the coding region, of which 247 were nsSNPs and 233 were synonymous SNPs (sSNPs). 888 frameshift. While 9.621 SNPs were in noncoding regions, of which 375 occurred in the 3’UTR, 131 in the 5’UTR region, and the rest (9115) were in the intronic region (Fig. S1). We chose nonsynonymous coding SNPs for our investigation (Fig. 1).

**Figure 1.**
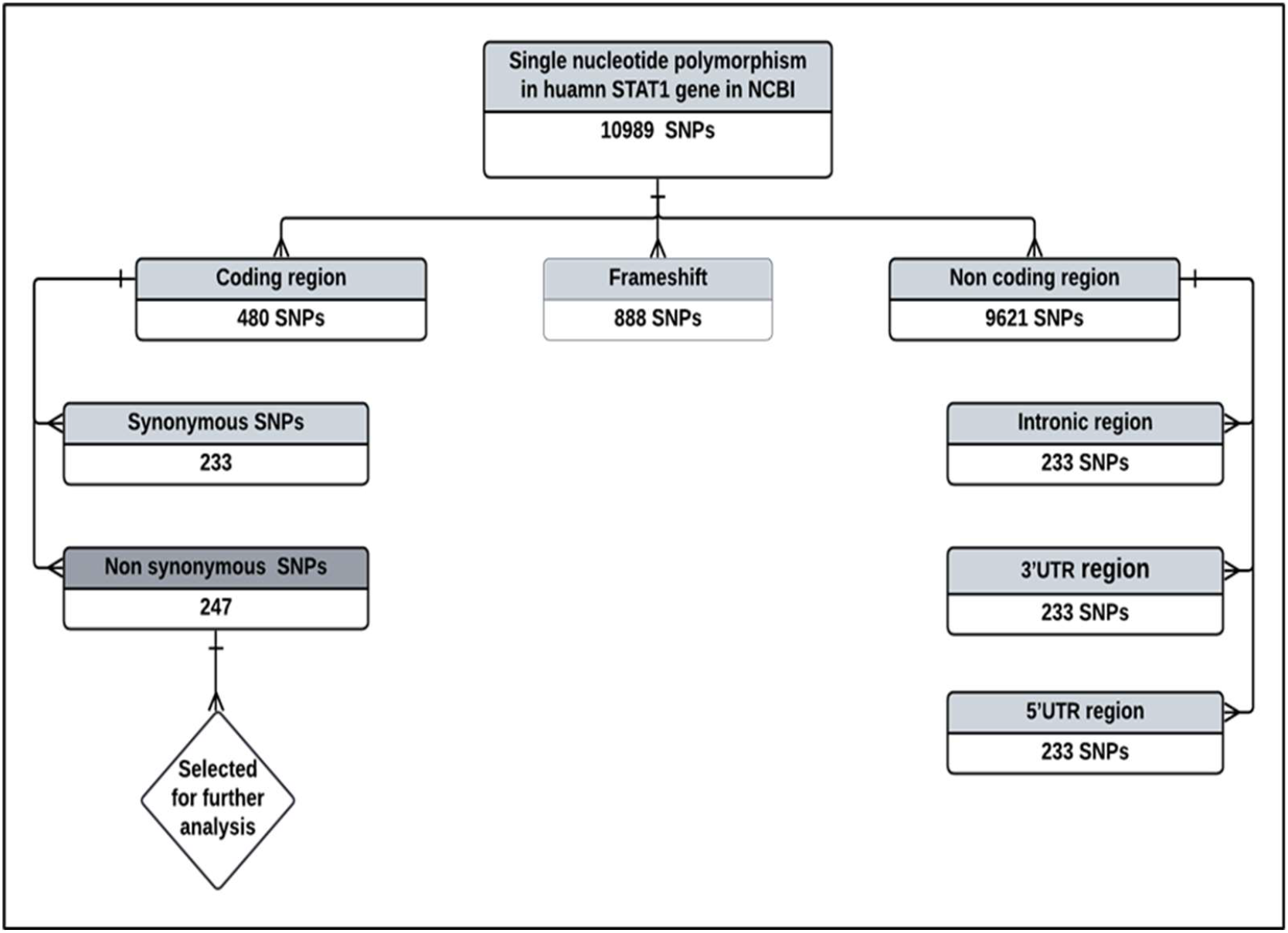
Selection of nsSNPs of *STAT1* gene.

An overview of the complete methodological approaches is summarized in the diagram (Fig. 2).

**Figure 2.**
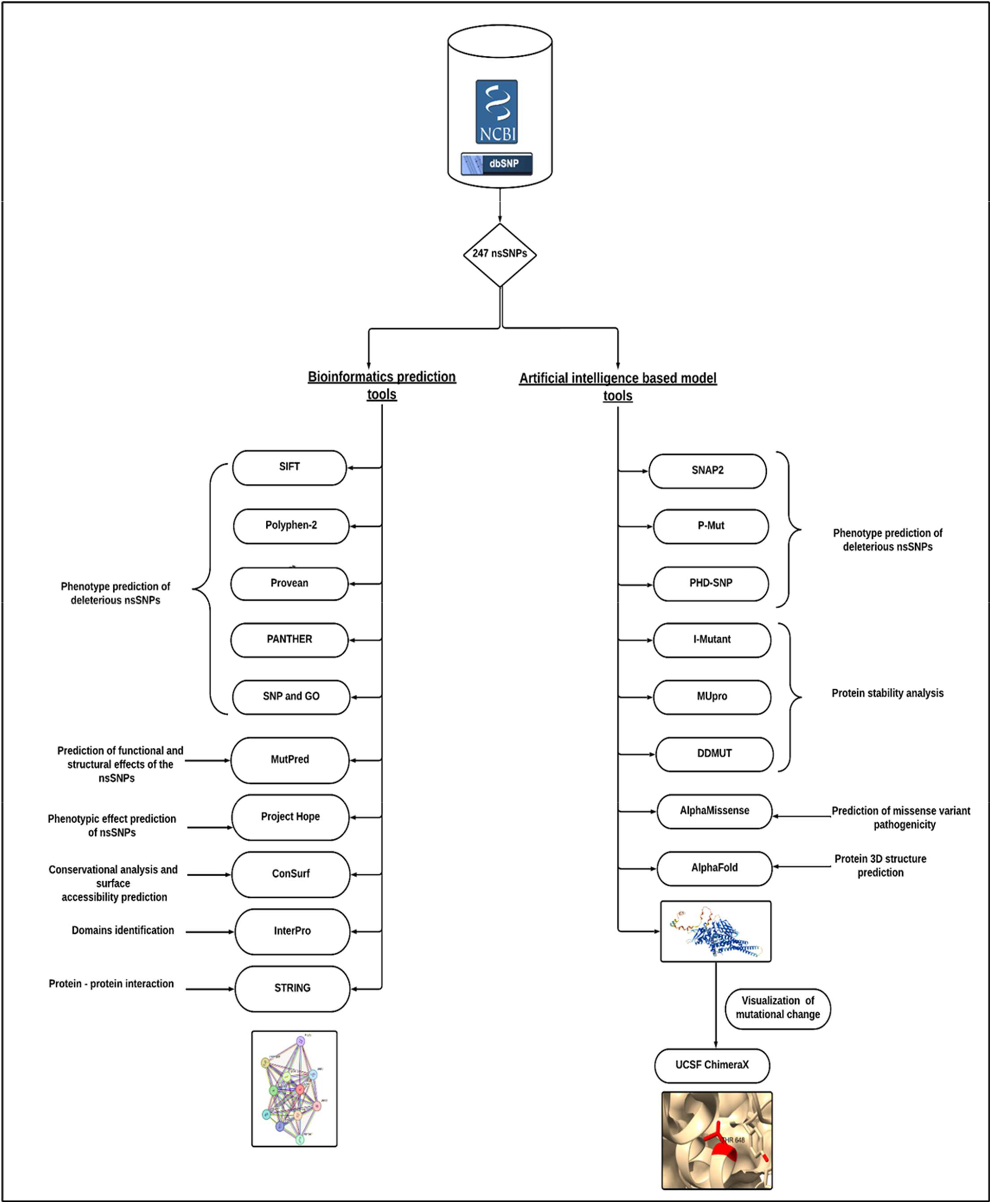
Workflow of the analysis.

### Single nucleotide polymorphism data retrieval

The data for the human STAT1 gene on chromosome 2q32 were gathered from the National Center for Biological Information (NCBI) website. While the SNP information (SNP ID) of the STAT1 gene was obtained from the NCBI dbSNP (http://www.ncbi.nlm.nih.gov/snp/), the protein ID and its sequence were extracted from Swiss-Prot databases with the accession number P42224. (http://expasy.org/).

### Identification of deleterious missense mutation

All 247 nsSNPs retrieved were subjected to pathogenicity prediction web servers. Sixty-four nsSNPs were found to be deleterious by SIFT and were further subjected to cross-checking by using three different tools (Poly-Phen-2, PROVEAN, and SNAP2). The shortlisted 33 nsSNPs were shown (Table 1).

**Table 1.** List of nsSNPs that were predicted to have deleterious effect by SIFT, PolyPhen-2, Provean and SNAP2

| SNP ID | Amino-acid change | SIFT Prediction | TI | Poly-Phen-2 Effect | Score | Provean Effect | Score | SNAP2 Prediction | Score |
| --- | --- | --- | --- | --- | --- | --- | --- | --- | --- |
| rs1173266737 | P728A | Deleterious | 0 | PB | 0.974 | Deleterious | -3.14 | E | 15 |
| rs1374373369 | D674V | Deleterious | 0.01 | PB | 0.989 | Deleterious | -7.327 | E | 47 |
| rs771679419 | Y668F | Deleterious | 0 | PS | 0.609 | Deleterious | -3.583 | E | 67 |
| rs759271255 | I648T | Deleterious | 0.01 | PB | 1 | Deleterious | -4.197 | E | 56 |
| rs752542806 | V642D | Deleterious | 0 | PB | 0.994 | Deleterious | -5.149 | E | 60 |
| rs1387961263 | L639F | Deleterious | 0 | PB | 1 | Deleterious | -3.489 | E | 7 |
| rs1209841496 | R602W | Deleterious | 0 | PB | 1 | Deleterious | -7.4 | E | 87 |
| rs137852678 | L600P | Deleterious | 0 | PB | 1 | Deleterious | -6.472 | E | 91 |
| rs1398307167 | P596Q | Deleterious | 0.01 | PS | 0.951 | Deleterious | -3.974 | E | 57 |
| rs1398307167 | P596L | Deleterious | 0.01 | PB | 0.966 | Deleterious | -5.853 | E | 58 |
| rs767475430 | I578N | Deleterious | 0 | PB | 0.1 | Deleterious | -6.392 | E | 80 |
| rs113988352 | I561T | Deleterious | 0 | PB | 0.981 | Deleterious | -4.143 | E | 38 |
| rs1803838 | P538L | Deleterious | 0.03 | PS | 0.588 | Deleterious | -3.097 | E | 10 |
| rs916580554 | W504C | Deleterious | 0 | PB | 0.999 | Deleterious | -11.937 | E | 48 |
| rs1185249247 | S503N | Deleterious | 0 | PS | 0.949 | Deleterious | -2.621 | E | 50 |
| rs866554932 | P481R | Deleterious | 0.04 | PB | 0.1 | Deleterious | -6.451 | E | 6 |
| rs935654762 | V455A | Deleterious | 0 | PB | 0.997 | Deleterious | -3.255 | E | 35 |
| rs527393923 | T450M | Deleterious | 0 | PB | 0.1 | Deleterious | -4.499 | E | 56 |
| rs760409880 | L448F | Deleterious | 0 | PS | 0.816 | Deleterious | -3.135 | E | 33 |
| rs776192196 | P326L | Deleterious | 0 | PB | 0.996 | Deleterious | -6.947 | E | 29 |
| rs763976174 | R304H | Deleterious | 0.02 | PS | 0.850 | Deleterious | -2.809 | E | 52 |
| rs751403509 | R304C | Deleterious | 0 | PB | 0.1 | Deleterious | -4.767 | E | 39 |
| rs779371351 | I248N | Deleterious | 0 | PB | 0.1 | Deleterious | -5.877 | E | 38 |
| rs779371351 | I248T | Deleterious | 0 | PB | 0.1 | Deleterious | -4.218 | E | 42 |
| rs1017740241 | C247Y | Deleterious | 0 | PB | 0.1 | Deleterious | -9.192 | E | 49 |
| rs763588438 | V149G | Deleterious | 0.01 | PB | 0.987 | Deleterious | -4.942 | E | 48 |
| rs1482374494 | A119T | Deleterious | 0 | PB | 0.1 | Deleterious | -3.113 | E | 26 |
| rs756147217 | P98S | Deleterious | 0 | PB | 0.1 | Deleterious | -5.885 | E | 48 |
| rs865962653 | S51L | Deleterious | 0.01 | PS | 0.883 | Deleterious | -4.4 | E | 34 |
| rs781389511 | A46T | Deleterious | 0 | PB | 0.1 | Deleterious | -2.543 | E | 1 |
| rs34255470 | I30T | Deleterious | 0.02 | PB | 0.1 | Deleterious | -3.391 | E | 17 |
| rs11549696 | P27T | Deleterious | 0 | PB | 0.1 | Deleterious | -6.503 | E | 64 |
| rs1233778383 | W4C | Deleterious | 0 | PB | 1 | Deleterious | -10.563 | E | 23 |

### MutPred prediction for functional and structural modifications

The shortlisted 9 nsSNPs were submitted to the MutPred server, along with the resultant probability scores and their P values (Table S2).The structural and functional alterations predicted include loss of disorder, catalytic residue, glycosylation, a gain of phosphorylation, solvent accessibility, ubiquitination, and molecular recognition features (MoRF) binding. According to these predictions, it can be stated that several nsSNPs might be the reason behind any possible structural and functional modifications of STAT1 protein.

### Prediction of change in STAT1 stability due to mutation using I-Mutant 3.0 server, MU pro, and DDMUT

Protein stability was analysed by I-mutant 3, MUpro, and DDMUT servers. The result revealed that six variants destabilized the STAT 1 residue, namely (I648T) rs759271255, (V642D) rs752542806, (R602W) rs1209841496, (L600P) rs137852678, (I578N) rs767475430, and (W504C) rs916580554 (Table S3).

### Pathogenicity prediction results

The analysis of STAT1 nsSNP prediction by Alpha Missense was performed, and we found that all the pathogenic nsSNPs that were predicted by the previous tools were also classified as pathogenic in Alpha Missense (Table S4). The heat map represented the mutations in STAT1 (Fig. S2).

### The conservational status and surface accessibility analysis of STAT1 protein

Highly conserved residues are most likely to be involved in proteins’ structural integrity and functions. The conservational profile was evaluated for the STAT1 protein. The ConSurf algorithm represented the structural and functional conservation levels of all the amino acid residues of the STAT1 protein (Fig. S3). Four SNPs (I648T, L600P, W504C, and I578N) are conserved and buried. L600P and I578Nare predicted structural residues (highly conserved and buried). V642D is buried, and R602W is predicted as functional residue (highly conserved and exposed) (Table S5).

### 3D structure prediction by AlphaFold and SNP visualization by ChimeraX

An individual residue confidence score (pLDDT) between 0 and 100 is generated by the AlphaFold algorithm. The majority of the 3D structural region corresponds to α-helical domains and has extremely high confidence (pLDDT>90). The remaining components of the model are depicted as unresolved loops with low (70>pLDDT>50) and extremely low (pLDDT>50) scores (Fig. S4). Alphafold produces a per residue confidence score (pLDDT) between 1-100. Regions with low pLDDT may be unstructured in isolation. ChimeraX is used to visualize the 3D structures of the wild-type amino acids in blue and the mutant residues in red (Fig. S5).

### The physical outcome of predicted SNPs

The impact of the generated damaging SNPs on the three-dimensional structure of STAT1 was examined using the HOPE server. The server predicted that all the mutated amino acids were different in size; 1 had a different charge, and 6 had different hydrophobicity (Table S6).

Loss of the interactions between the wild-type amino acid and other amino acids in the protein and/or development of new interactions or bonds between the mutant residue of the protein and the other amino acids in the protein were predicted by DDMUT (Fig. S6).

### Domain identification of the STAT1 protein by the InterPro server

The InterPro tool predicted the domain regions of the STAT1 protein. It is mainly reported that the STAT1, SH2-domain (a phosphotyrosine binding pocket) at position (557–707), STAT transcription factor, DNA binding domain at (323–458), and STAT1_TAZ2-binding domain (715-739) are conserved sites. Src homology 2 (SH2) domain profile (573-670), SH2 domain (578-638). STAT1 transcription factor, all alpha domain (144–305), and STAT transcription factor, protein interaction (2–12) (Table S7).

### STAT1-protein interaction

Analysis of protein-protein interaction The STRING network revealed that STAT1 interacts with 10 proteins, which include other proteins of the same STAT family (STAT2, and STAT3), proteins of the JAK family (JAK1, and JAK2), IFR1, IFR9, IFNGR1, CREBBP, KBNA1, and PIAS1 (Figure. S7).

## Discussion

Bioinformatics tools are used to investigate the influence of missense SNPs on STAT1 protein focusing on their structural and functional consequences. By applying various artificial intelligence-based models and bioinformatics tools, we identified six pathogenic variants (R602W, I648T, V642D, L600P, I578N, and W504C) within the STAT1 gene that are predicted to significantly disrupt the protein stability and function. These findings underscore the potential of this approach to pinpoint Pathogenic SNPs, providing a time and cost-effective alternative to experimental approaches. Such an advantage will encourage researchers to utilize computational methods to prioritize the most harmful genetic variants for further experimental investigation, specifically in genes like STAT1 which plays a crucial role in immune response function and disease susceptibility.

The objective of this study is to define the structural and functional characterization of the most pathogenic variations of the *STAT1* gene. We evaluated the functional and pathogenic sequences of missense SNPs of the human *STAT1* gene utilizing thirteen diverse in silico prediction tools (SIFT, PolyPhen2, PROVEAN, PANTHER, P MUT, PhD-SNP, SNPs&GO, SNAP2, MutPred2, I-Mutant, MUpro,, DDMUT, and alpha-missense).

In-silico prediction analysis identified six variants (I648T, V642D, R602W, L600P, I578N, and W504C) to be considered pathogenic and deleterious. These mutations have a major impact on the protein’s physicochemical characteristics, such as its size and charge hydrophobicity, which ultimately affect the protein’s stability and function and may have an impact on disease. Furthermore, we assessed the effect of missense SNPs on the stability of the STAT1 structure utilizing two stability predictions.

Algorithms: I-Mutant3, MUpro, and DDMUT. All the variants revealed a reduction in stability by the three stability prediction tools (I-Mutant3, MUpro, and DDMUT). In general, we assumed that all missense SNPs in the *STAT1* gene were highly unstable in their protein structures, so they were selected for further structural bioinformatics analysis utilizing various tools to explore the Consequences of tentatively destructive missense SNPs on STAT1 protein function. To evaluate the conservation profile of STAT1 protein, we used the ConSurf algorithm to represent the structural and functional conservation levels of all the amino acid residues of STAT1 protein. The ConSurf analysis revealed that the variant in position 602 is functional residue in a highly conserved and exposed position. Structural residues in highly conserved and buried positions were identified in positions 600 and 578. The identified variants were found in a highly conserved region; this finding suggests that they might be involved in modifications of molecular mechanisms such as bond gain or loss.STAT1 GOF mutations with CMC were first described in 2001 and 2011, respectively; later, studies confirmed that STAT1-GOF mutations cause immunodeficiency and immune dysregulation, with a wide clinical spectrum ^[19]^.

Among the six SNPs identified to be linked to STAT1 gene mutations in this study, some of these SNPs have been associated with diseases in previous studies, while others were projected to be so in this study using various computational tools. Population genetics and clinical studies are crucial to verifying the results of such research, even though utilizing computational techniques to analyse the impact of the SNPs may aid in identifying disease-related SNPs.

One mutation, namely L600P, has already been previously reported as a mutation in the STAT1 gene in an infant who died of a viral-like illness associated with complete STAT1 deficiency and carried a homozygous nucleotide substitution (T→C) in exon 20, resulting in the substitution of proline for leucine at amino-acid position 600. This mutation was found to be pathogenic using all the bioinformatics tools ^[20]^; I648T, V642D, R602W, I578N, and W504C were not reported previously. Three mutations ^[21]^, namely L706S (rsRCV000009610), Q463H (VAR_065817), and E320Q (VAR_065816) have been reported as mutations in the *STAT1* gene. The two previously reported types of autosomal dominant (AD) Mendelian susceptibility to mycobacterial disease (AD-MSMD) causing STAT1 mutations are located in the tail segment domain (p.L706S) or the DNA-binding domain (p.E320Q and p.Q463H)^[21]^. These mutations were not available in the dbSNP database.

two other SNPs (K637E) and (K673R) affecting the SH2 domain, which has been previously reported in two cases with AD-STAT1 deficiency in two unrelated patients from Japan and Saudi Arabia, were also not available in the dbSNPs database at the time of the analysis.^[21]^ two mutations were linked to chronic mucocutaneous candidiasis (T437I) and (Q271P); the latter occurred within a specific pocket of the STAT1 coiled-coil domain, near residues essential for dephosphorylation, and was identified in a German patient who presented at 1 year of age with autosomal dominant chronic mucocutaneous candidiasis, showed signs of thyroid autoimmunity, and died at age 41 from squamous cell carcinoma^[22,23]^. These mutations were not available in the dbSNP database.

The Ala267Val variant in STAT1 has been reported in >10 individuals with chronic mucocutaneous candidiasis (CMC) and segregated with disease in 16 individuals from 9 families ^[24]^. This important mutation was not present in the dbSNP database.

Interestingly, nsSNPs in the *STAT1* gene will ultimately affect and may disturb the normal functioning of other interacting genes. As our study was in detail, it provides all the information and analysis that are needed for the identification of the most damaging nsSNPs. Like ours, there are certain limitations in every study. Our study is based on computer tools and web servers, which are based on mathematical and statistical algorithms. Therefore, to confirm these results, experimental investigation is necessary to confirm their functional effects and clinical relevance. Our study provides insight into nsSNPs of the *STAT1* gene, its protein 3D structure, and its gene-gene interaction with other genes, which might be helpful in future studies of STAT1 to better understand its role in immunity and all related diseases. Additionally, including population genetics data may yield insights into the incidence and distribution of these polymorphisms among other groups, facilitating the development of individualized treatment approaches.

## Materials and Method

### Single nucleotide polymorphism data retrieval

The data for the human STAT1 gene were gathered from the National Center for Biological Information (NCBI) website (https://www.ncbi.nlm.nih.gov/). While the SNP information (SNP ID) of the STAT1 gene was obtained from the NCBI dbSNP (http://www.ncbi.nlm.nih.gov/snp/), the protein ID and its sequence were extracted from Swiss-Prot databases with the accession number P42224. (http://expasy.org/)^[25]^.

### Phenotype prediction of deleterious ns SNPs

Prediction of the deleterious nsSNPs was performed by using different eight tools. Sorting Intolerant from Tolerant (SIFT) (http://sift.bii.a-star.edu.sg/) ^[26]^. Polyphen-2 (http://genetics.bwh.harvard.edu/pph2/)^[27]^. Provean (https://www.jcvi.org/research/provean/.)^[28]^. SNAP2 (https://rostlab.org/services/snap2web/.)^[29]^. PHD-SNP (https://snps.biofold.org/phd-snp/phd-snp.html.)^[30]^.SNP and GO (https://snps-and-go.biocomp.unibo.it/snps-and-go/)^[31]^. P-Mut (http://mmb.irbbarcelona.org/PMut.)^[32]^, and Protein Analysis through Evolutionary Relationships (PANTHER) (http://pantherdb.org/) ^[33]^.

SIFT predicts if the replacement of an amino acid alters protein function. We downloaded nsSNP IDs from the online databases of NCBI and then uploaded them to SIFT. Results were documented as damaging (deleterious) or benign (tolerated), depending on the cutoff value of 0.05, as values less than or equal to (0.0–0.04) were predicted to be damaging or intolerant, while (0.05_1) is benign or tolerated. Polyphen-2 analyzes multiple sequence alignments and the protein’s three-dimensional structure, then predicts the possible impact of amino acid substitutions on the stability and function of human proteins using structural and comparative evolutionary considerations. Prediction outcomes are classified as probably damaging, possibly damaging, or benign based on the PSIC value, which ranges from 0 to 1. Values near zero are regarded as benign, while values near 1 are regarded as probably damaging. Provean is a software tool that predicts whether an amino acid substitution or indel has an impact on the biological function of a protein. Variants with a score equal to or below -2.5 are considered “deleterious,” while variants with a score above -2.5 are considered “neutral.” SNAP2 is a trained classifier that uses the “neural network” machine learning tool to predict the functional effects of mutations by utilizing several sequence and variant properties to discriminate between effect and neutral variants/non-synonymous SNPs. PHD-SNP uses a support vector machine (SVM)-based method trained to determine disease-associated nsSNPs using sequence information. The related mutation is predicted to be disease-related (disease) or a neutral polymorphism. SNP and GO is a server for the prediction of single-point protein mutations likely to be involved in the development of diseases in humans. P-Mut is a web-based tool for the annotation of pathological variants on proteins. It allows fast and accurate prediction of the pathological properties of single-point amino acid mutations based on the use of a neural network. PANTHER (Protein Analysis through Evolutionary Relationships) uses a position-specific evolutionary preservation (PSEP) score to measure the length of time (in millions of years) with < 200 my ‘probably benign’, < 450 my ‘possibly damaging’, and 450 my ‘probably damaging.

### Predicting functional and structural effects of the nsSNP

MutPred v1.2 (http://mutpred.mutdb.org/), ^[34]^ was used for sorting disease-associated or neutral amino acid substitutions in humans. It is a web-based application tool that efficiently screens amino acid substitutions and predicts the molecular base of the disease.

### Protein stability analysis of predicted STAT1 nsSNPs

I-Mutant 3.0 is a neural network-based tool for routinely analysing protein stability and change while taking single-site mutations into consideration ^[35]^. The FASTA sequence of proteins retrieved from UniProt is used as an input to predict the mutational effect on protein stability. It is available at https://gpcr2.biocomp.unibo.it/cgi/predictors/I-Mutant3.0/I-Mutant3.0.cgi.

MUpro, a group of machine learning methods, has been developed for predicting the effects of single amino acid substitutions on protein stability ^[36]^. It uses both support vector machines and neural networks; the output is either increased or decreased stability ^[36]^. MUpro also interprets the result based on Gibbs free energy (ΔΔG) with a confidence score between − 1 and 11. MUpro is available at http://mupro.proteomics.ics.uci.edu.

DDMUT (https://biosig.lab.uq.edu.au/ddmut/.) Is a fast and accurate network using deep learning models to predict changes in Gibbs free energy (ΔΔG) upon single and multiple point mutations ^[37]^. DDMut achieved Pearson’s correlation of up to 0.70 (RMSE: 1.37 kcal/mol) on predicting single-point mutations on cross-validation and 0.74 (RMSE: 1.67 kcal/mol) on multiple mutations.

### Prediction of missense variant Pathogenicity

Alpha Missense is an adaptation of alphafold fine-tuned on human and primate variant population frequency databases to predict missense variant pathogenicity. It works by combining structural context and evolutionary conservation. This model achieves state-of-the-art results across a wide range of genetic and experimental benchmarks, all without explicitly training on such data.^[38]^

### 3D structure prediction and visualization

The 3D structure was predicted using an artificial intelligence system, AlphaFold (https://alphafold.ebi.ac.uk). AlphaFold is an AI system developed by Google DeepMind that predicts a protein’s 3D structure from its amino acid sequence. It can predict protein structures computationally with high accuracy ^[39]^. The UniProt sequence of the STAT1 protein was used as an input to get the alpha-fold model.

UCSF ChimeraX is a robust application that enables interactive viewing and analysis of various molecular structures and related data, including density maps, sequence alignments, and supramolecular assemblies ^[40]^. It allows the mapping and visualization of amino acid substitutions. Chimera X is found at https://www.rbvi.ucsf.edu/chimerax/.

### Phenotypic effects prediction

Project Hope (version 1.0) is an online web server used to analyse the structural and conformational variations that have resulted from single amino acid substitutions ^[41]^. Protein sequences, wild-type Amino acids and mutants were selected. The results provided describe the change in the physiochemical properties of the amino acid in the given SNPs. It is available at (http://www.cmbi.ru.nl/hope).

DDMUT can also detect changes in the biological interactions between wild-type amino acids and neighbourhood residues in comparison with mutant residues ^[37]^.

### Conservational analysis and surface accessibility prediction of STAT1

The ConSurf bioinformatics tool (https://consurf.tau.ac.il) was used to study the evolutionary conservation of nsSNP positions in a protein sequence ^[42]^. In this tool, phylogenetic relations were analysed between homologous proteins. The FASTA sequence of the STAT1 protein was submitted to the server, and the highly conserved residues were screened out. Identification of nsSNPs in STAT1 protein domains

The FASTA sequence of the STAT1 protein was submitted to the InterPro server (https://www.ebi.ac.uk/interpro). It predicts protein families and conserved domains, and then the positions of nsSNPs were manually pinpointed within these domains ^[43]^.

### Prediction of protein-protein interactions

A pre-computed database, STRING (https://string-db.org/), is used to determine protein-protein interactions to understand the function, structure, molecular action, and regulation of the protein ^[44]^. The protein sequence was used as an input query. **Conclusions**

Taken together, the study offers a first-time and extensive *in silico* analysis of missense SNPs in the STAT1 activity and the potential induced pathogenicity, revealing interesting pathogenic variants that may contribute to immune dysregulation. While bioinformatics and integrated artificial intelligence-based model tools offer valuable insight, further wet lab experimental validation is essential to confirm the molecular and clinical relevance of these findings, which will ultimately improve our understanding of *STAT1*-related diseases and inform population-based therapeutic strategies.

## Acknowledgements

We acknowledge and thank all participants for their cooperation and scientific contributions.

## Author contributions

Conceptualization, E.K, L.A.K, M.A.; Data curation, E.K, A.A, M.A.; Funding acquisition, E.K.; Investigation, E.K, M.A, L.A.K and A.A.; Methodology, E.K, M.A.; Software, E.K and L.A.K.; Supervision, AA.; Validation, E.K, M.A.; Writing—original draft, E.K, L.A.K, M.A, and AA.; Writing—review & editing, E.K.,M.A., A.A.

## Corresponding authors

Correspondence to Ebtihal Kamal

## Ethics declarations

No human or animal samples were used in this study

## Competing interests

The authors declare no competing interests.

Supplementary Information: online

