## Supplementary figures and images for "Integrating Artificial Intelligence and Bioinformatics Methods to Identify Disruptive STAT1 Variants Impacting Protein Stability and Function"

### supplementary figure 1

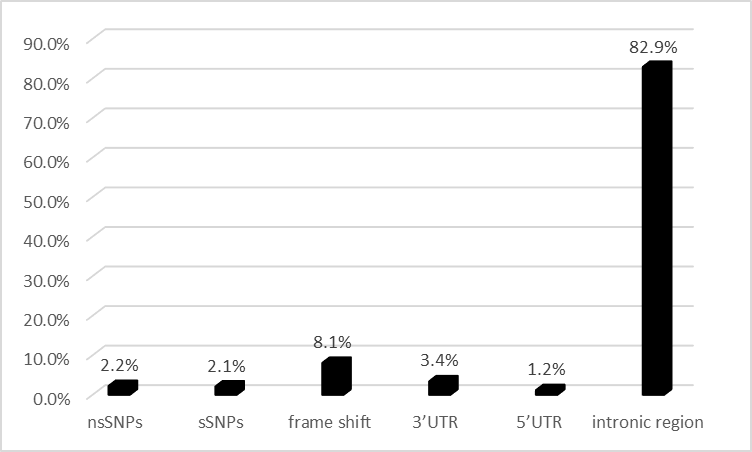

Figure S1. Shows the percentages of the SNPs in *STAT1* gene (nsSNPs: 1.8%; 3’UTR SNPs: 3.4%; 5’UTR SNPs: 1.2%; Other SNPs: 93.6%)

### supplementary figure 2

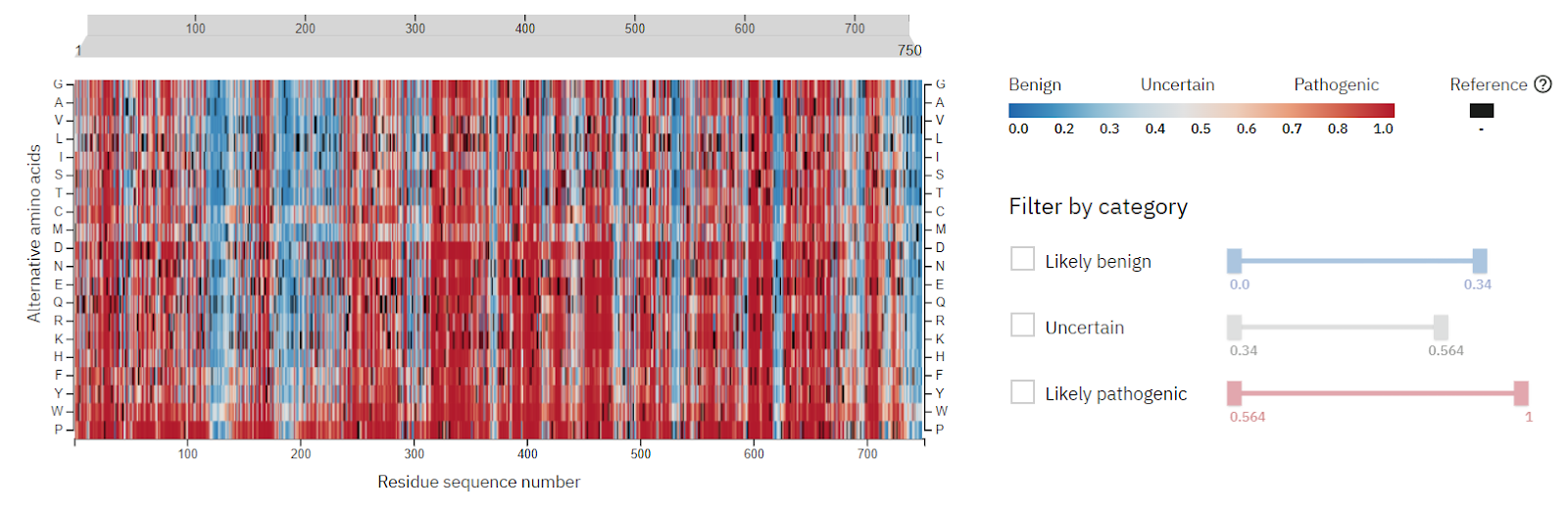

Figure S2. Heat map generated by alpha-missense, shows the variations in *STAT1* gene.

### supplementary figure 3

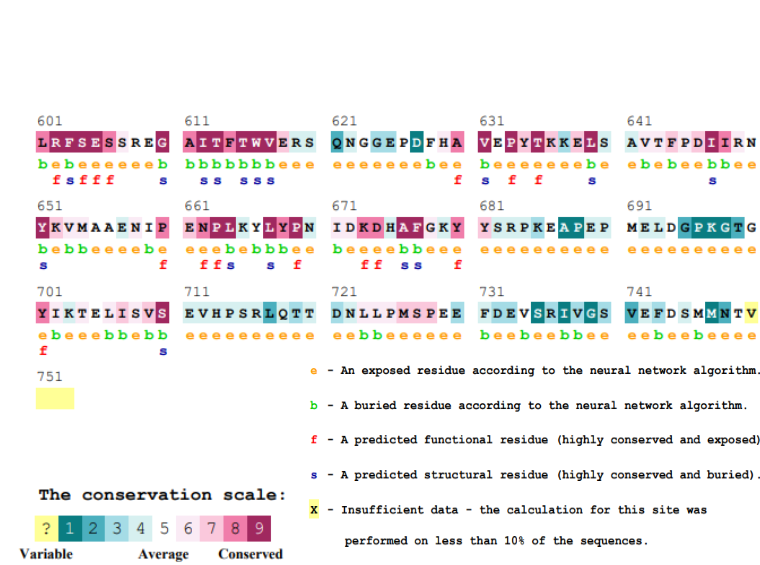

Figure S3. Conservation profile of amino acids in STAT1protein

### supplementary figure 4

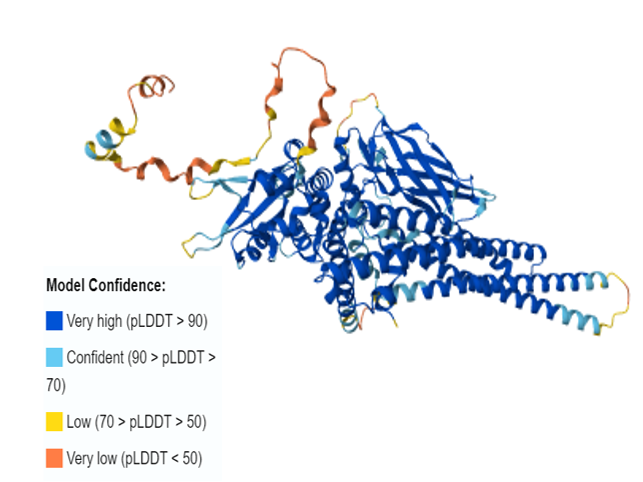

Figure S4. Protein 2D structure of human STAT1predicted by AlphaFold2

### supplementary figure 7

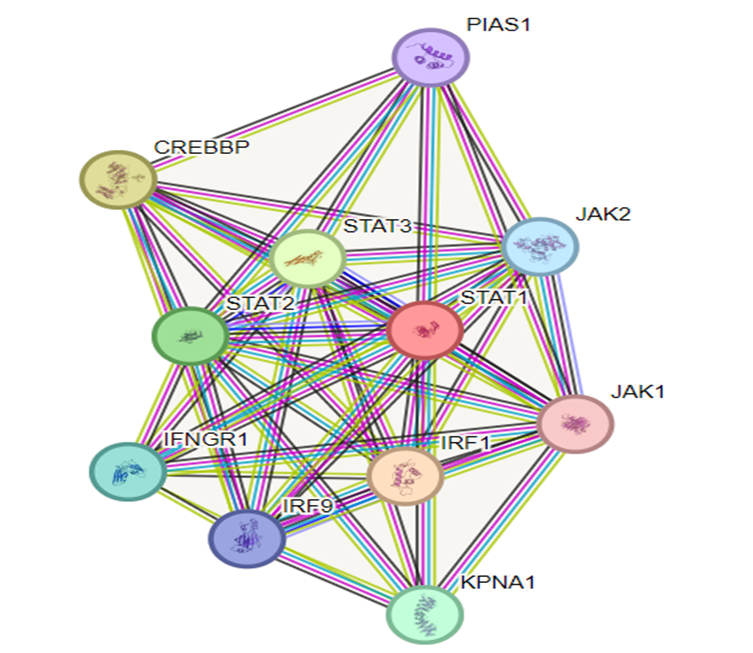

Figure S7. STAT1-protein interactions
