## supplementary figure 5 for "Integrating Artificial Intelligence and Bioinformatics Methods to Identify Disruptive STAT1 Variants Impacting Protein Stability and Function"

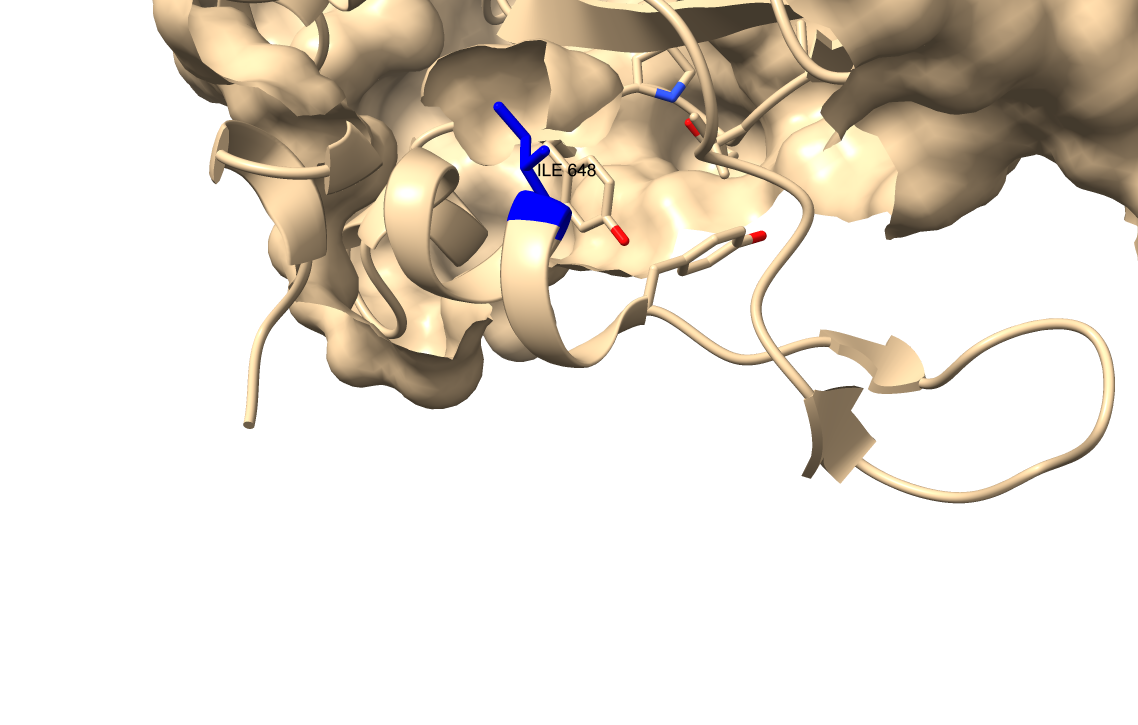

Wild type amino acid at position 648.

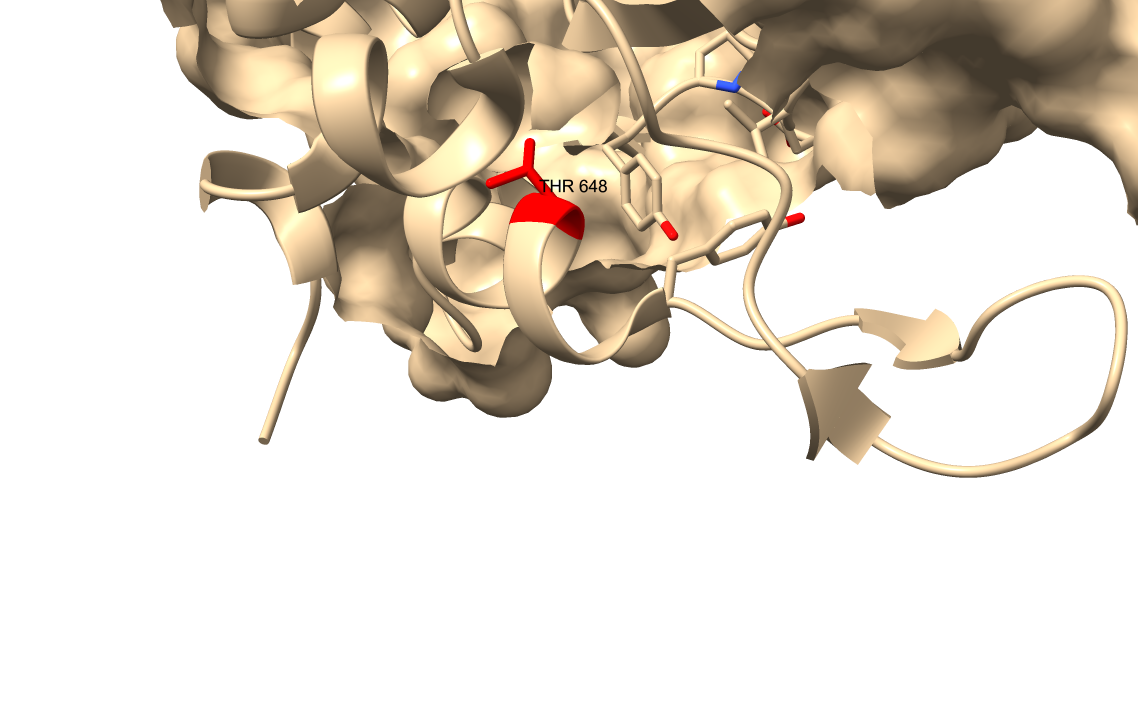

Mutant residue is at position 648.

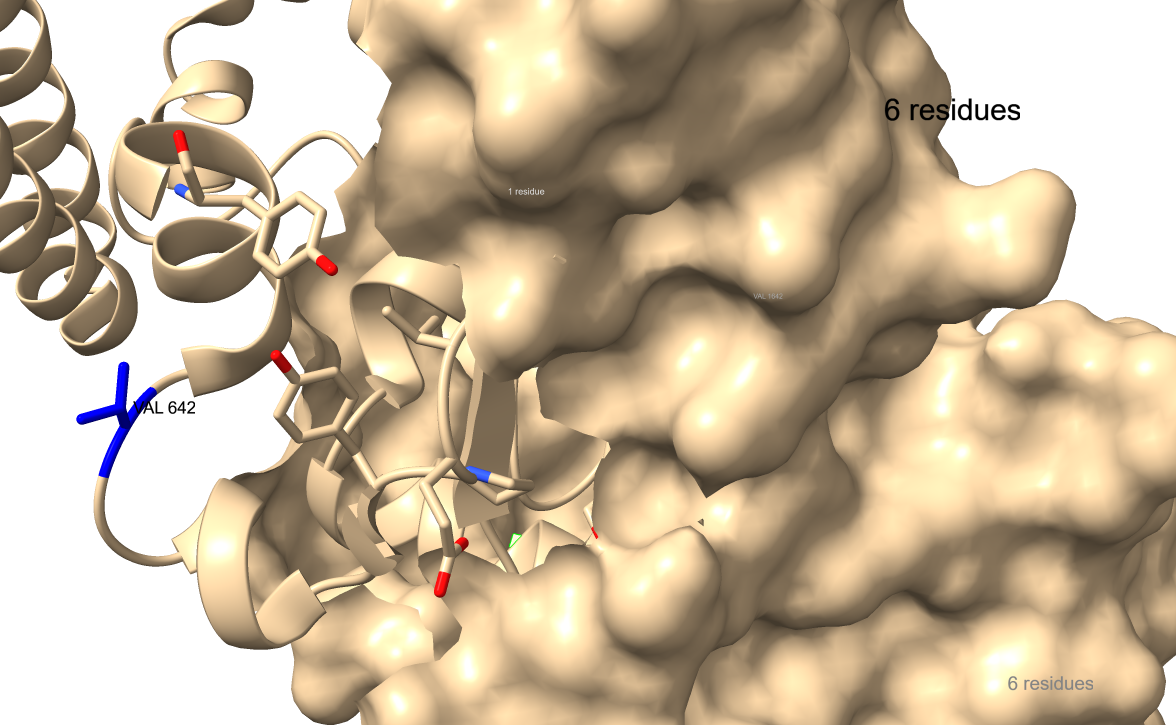

Wild-type amino acid at position 642.

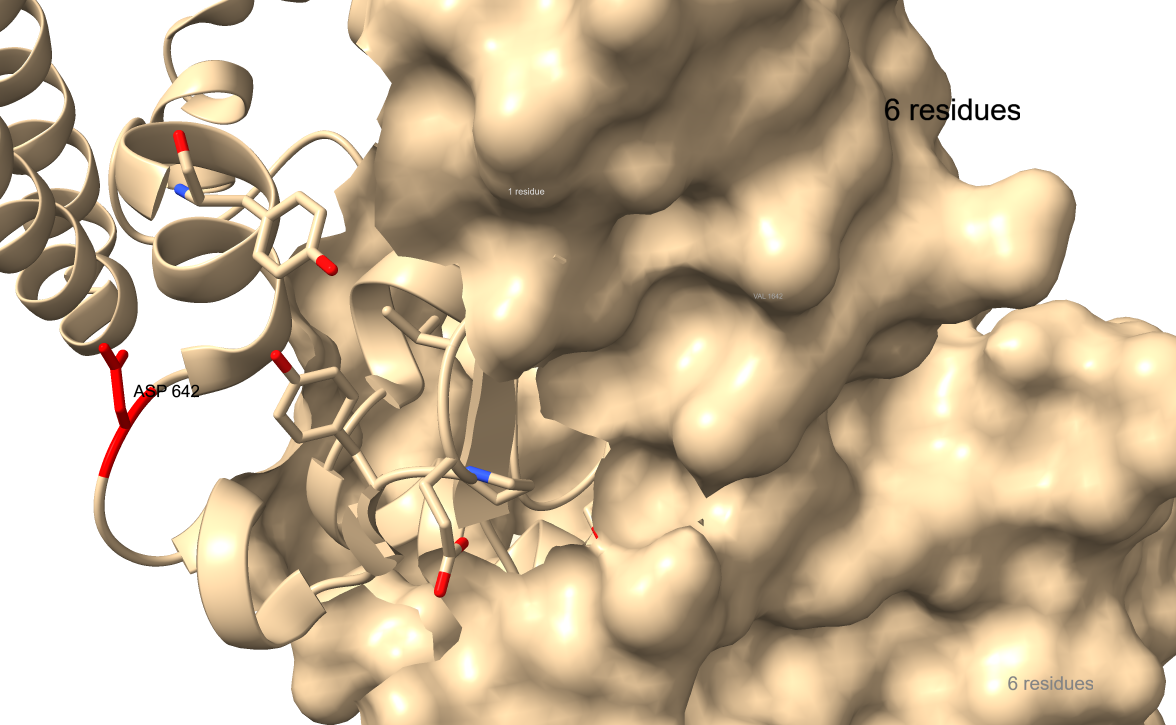

Mutant residue is at position 642.

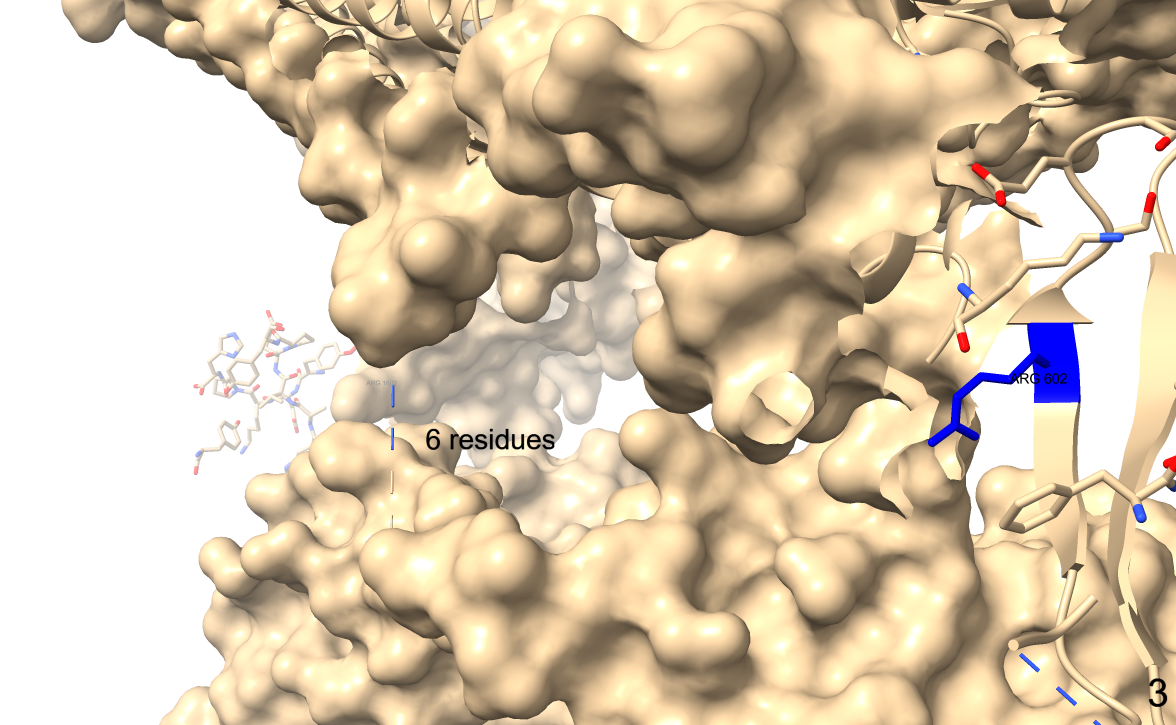

Wild type amino acid at position 602.

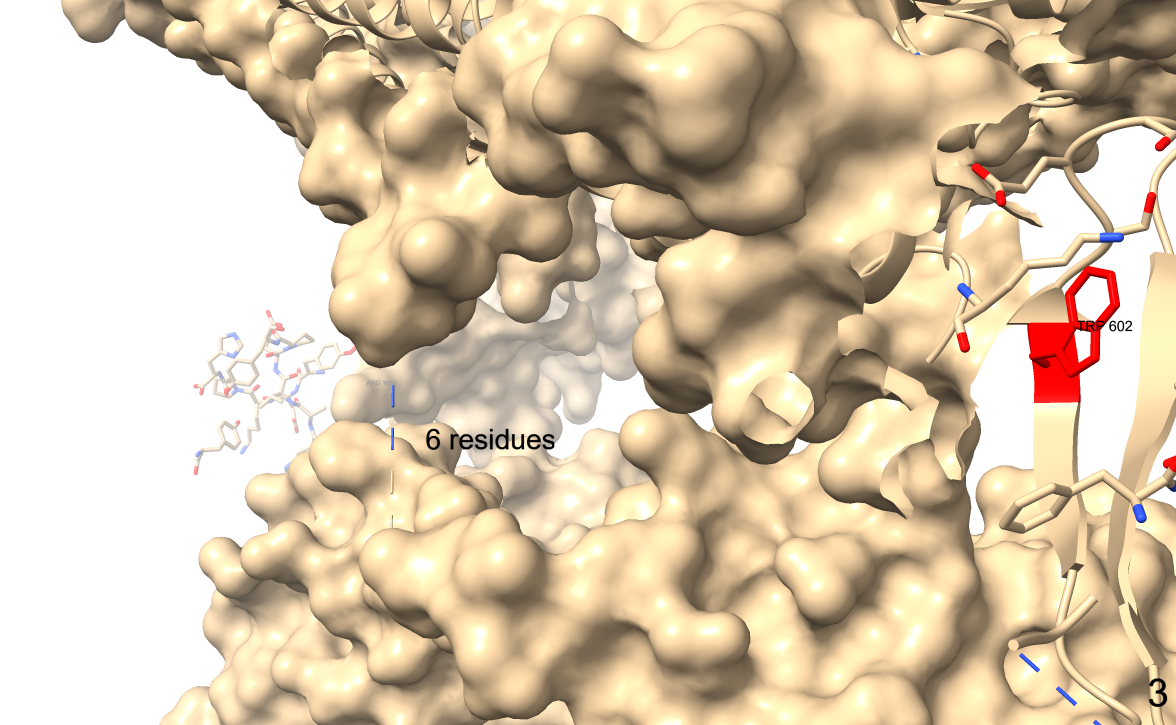

Mutant residue at position 602.

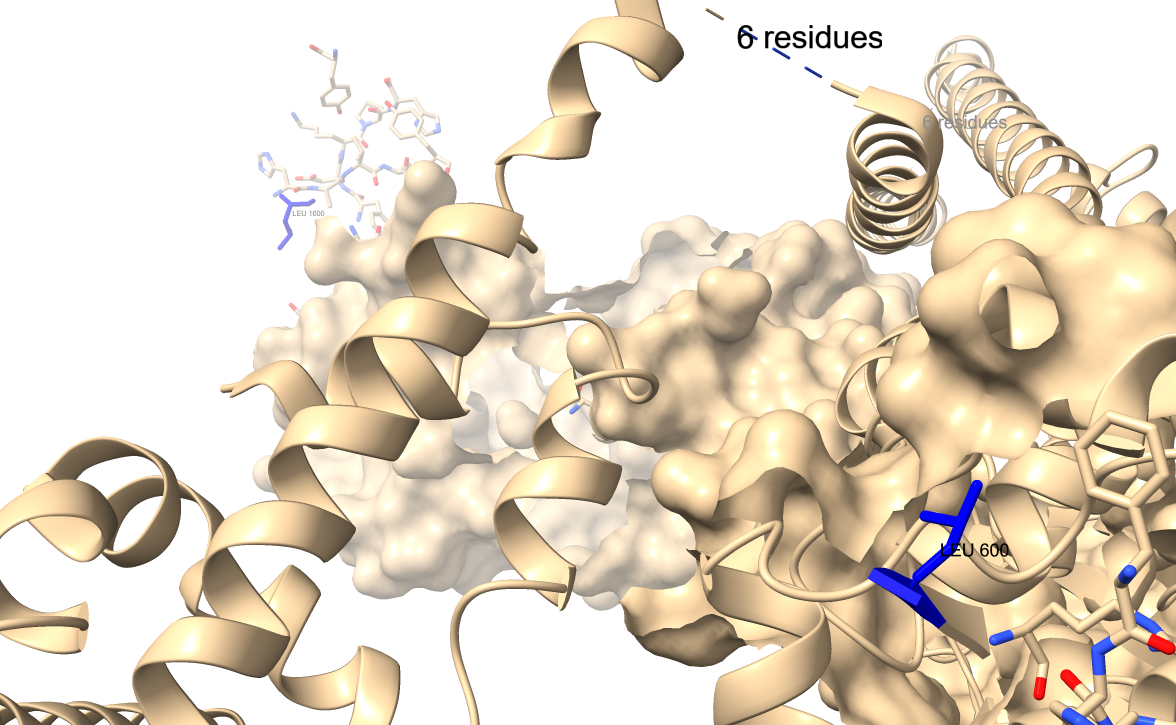

Wild type amino acid at position 600.

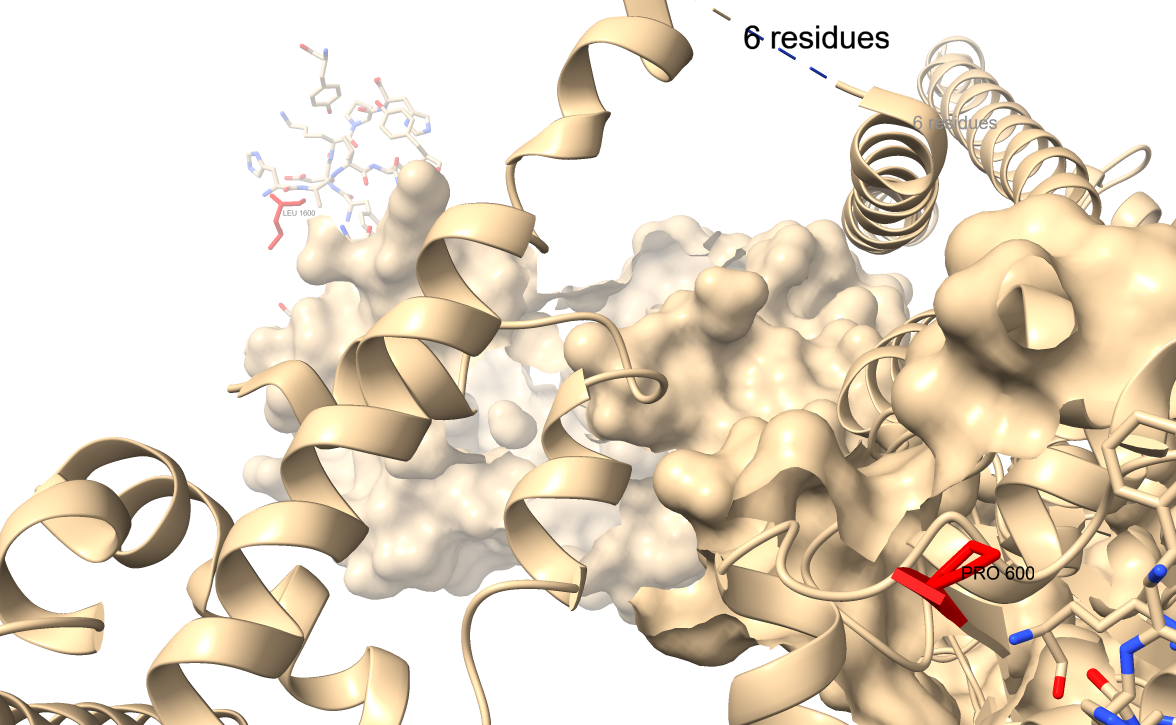

Mutant residue at position 600.

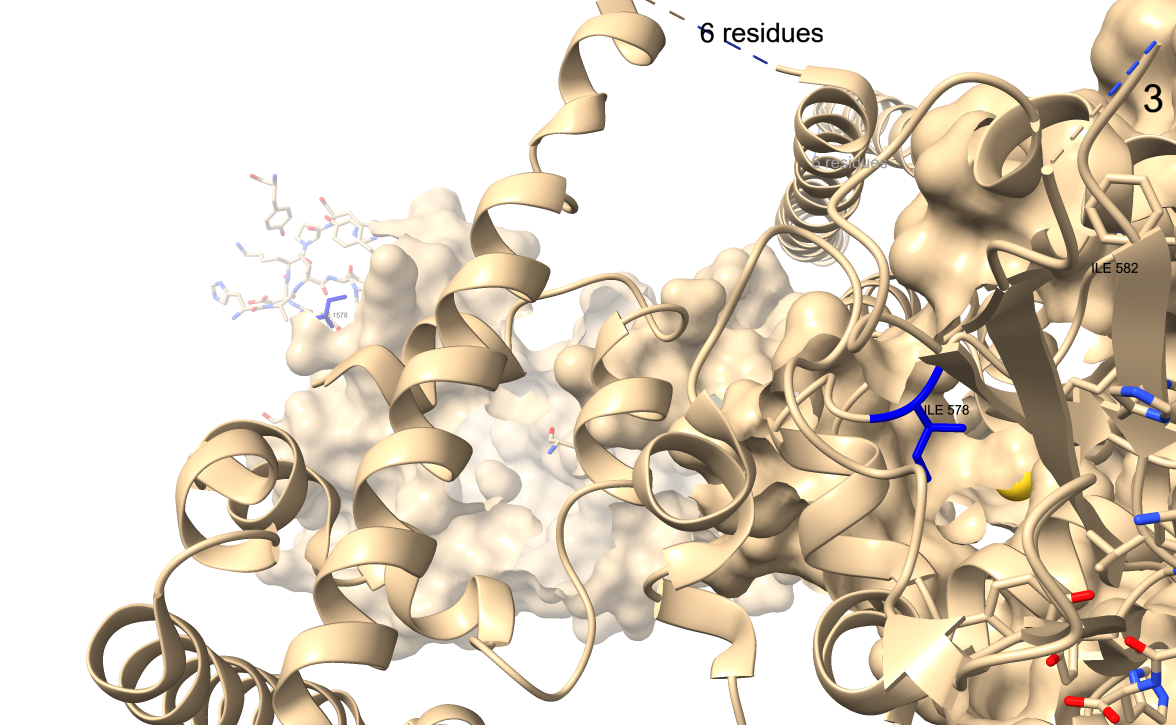

Wild-type amino acid at position 578.

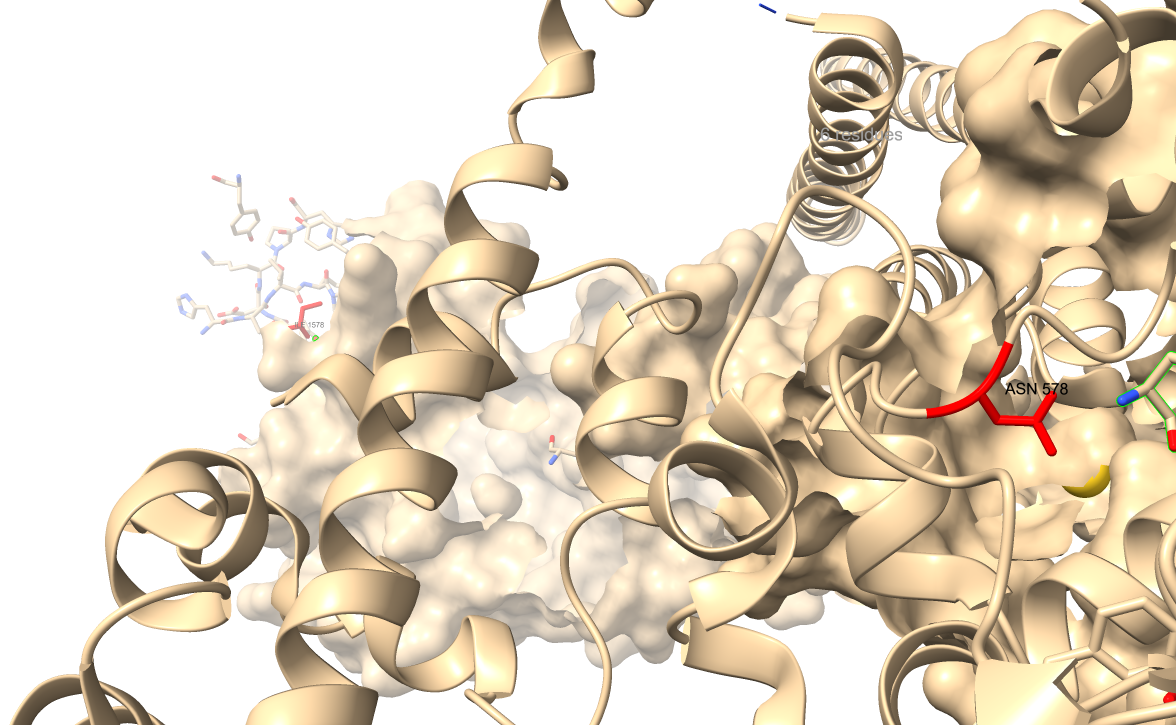

Mutant residue at position 578.

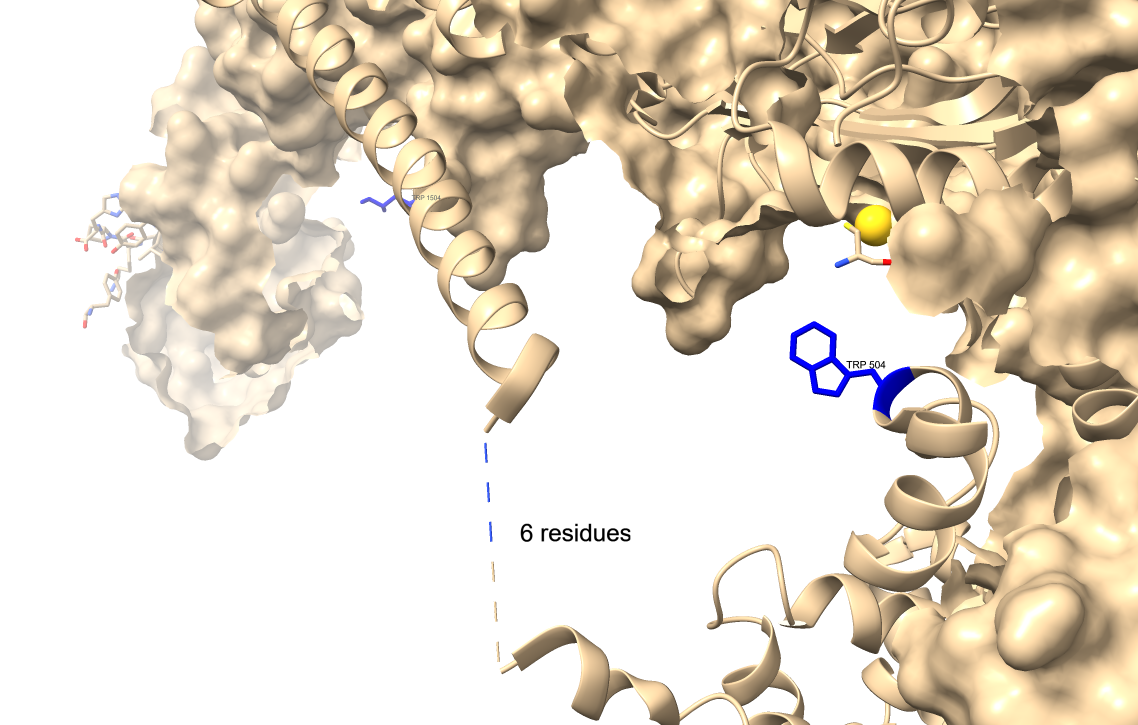

Wild type amino acid at position 504.

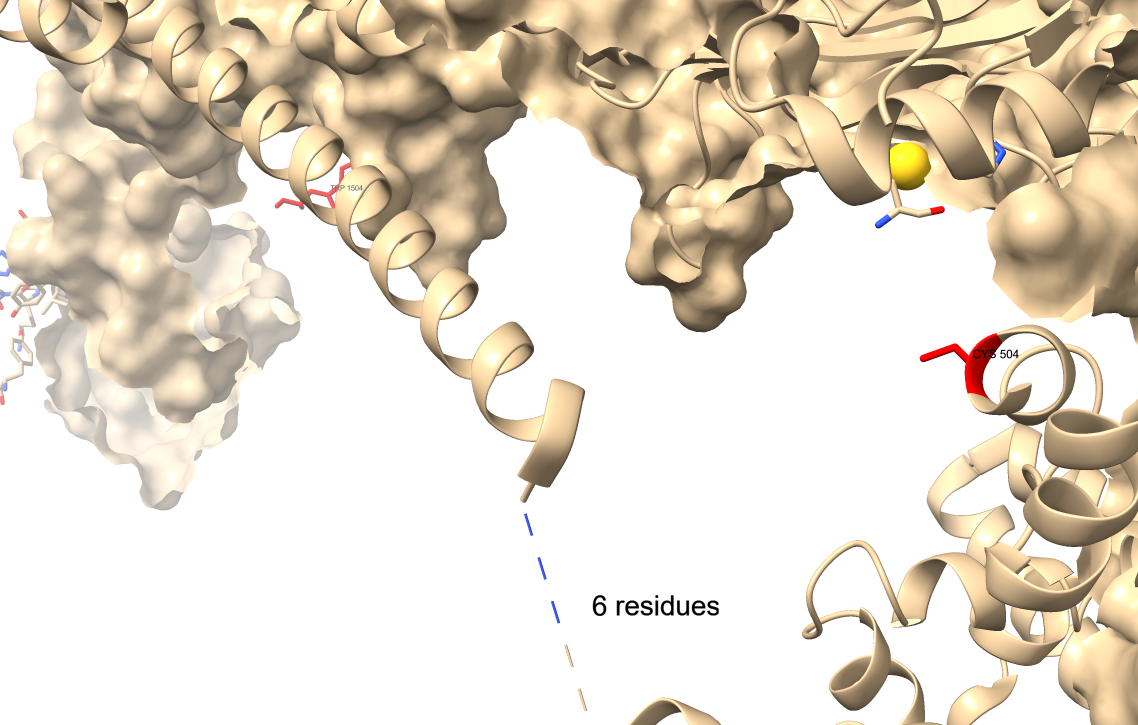

Mutant residue at position 504.

Figure S5. Effect of the six most deleterious nsSNPs on the STAT1 protein structure. ChimeraX software was used to visualize the 3D structure.
