## supplementary figure 6 for "Integrating Artificial Intelligence and Bioinformatics Methods to Identify Disruptive STAT1 Variants Impacting Protein Stability and Function"

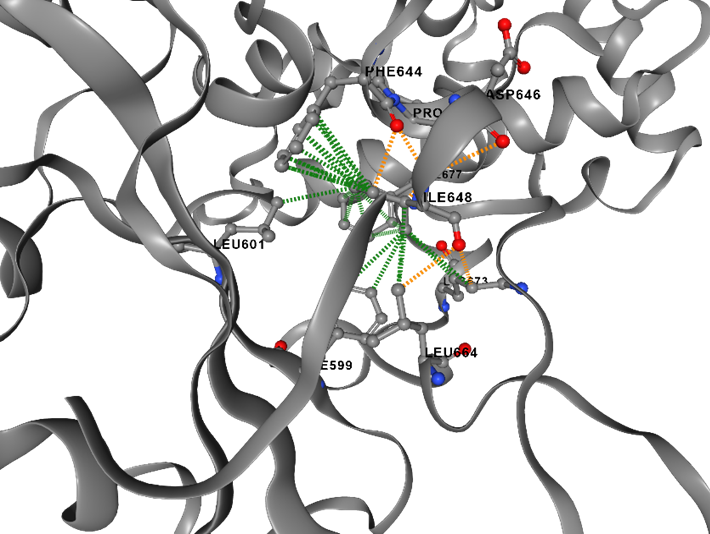

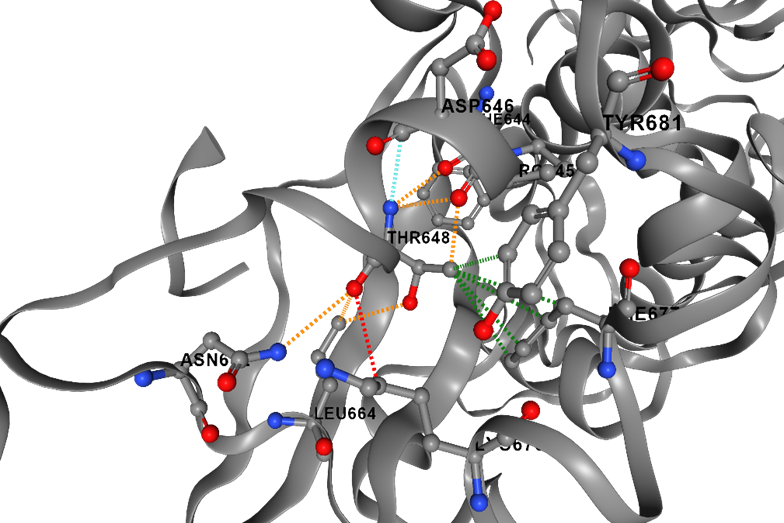

(A)I648T - wild type	(B) I648T, mutant type

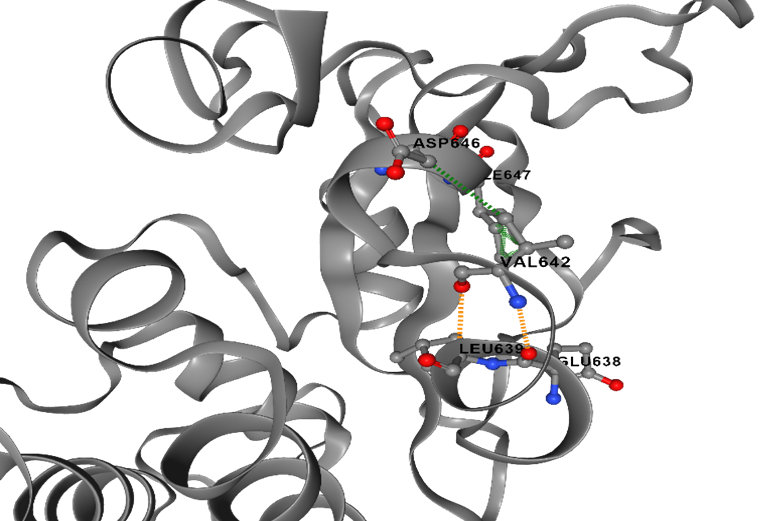

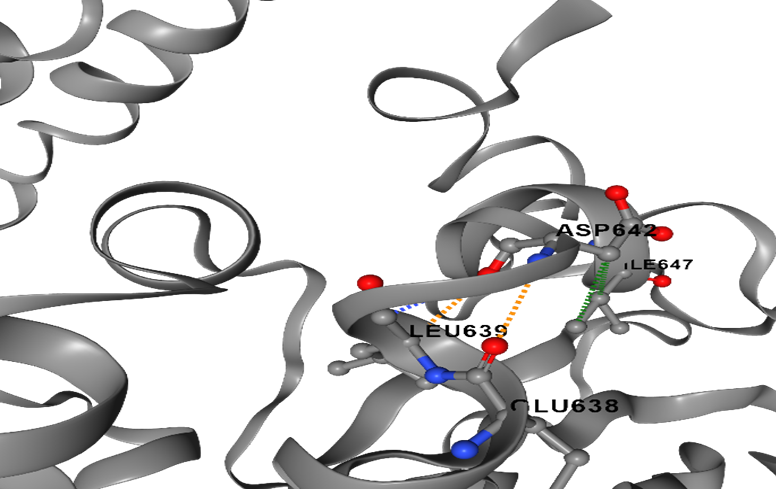

(A)V642D - wild type	(B) V642D, mutant residue

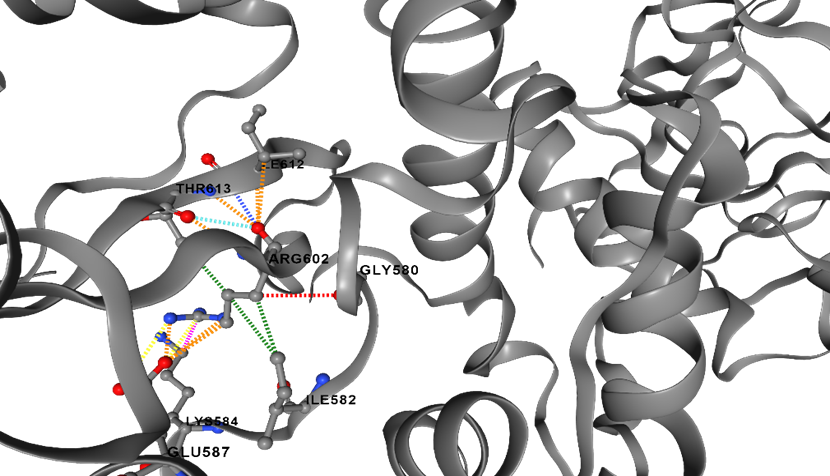

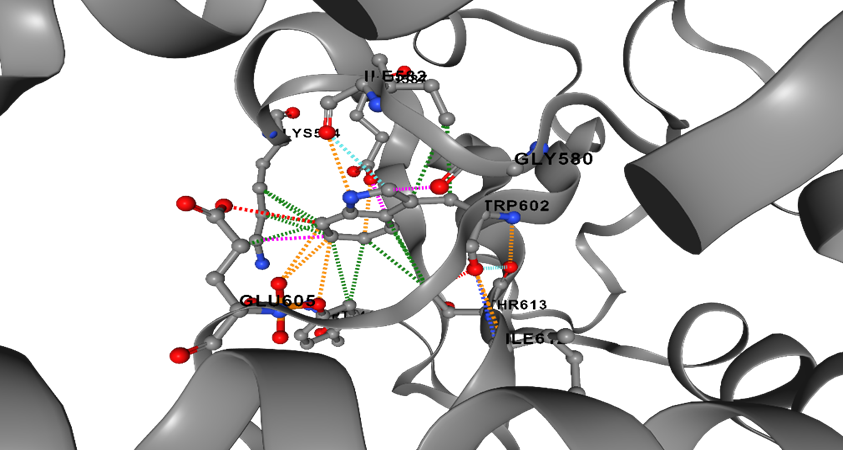

(A)R602W - wild type	(B) R602W, mutant residue

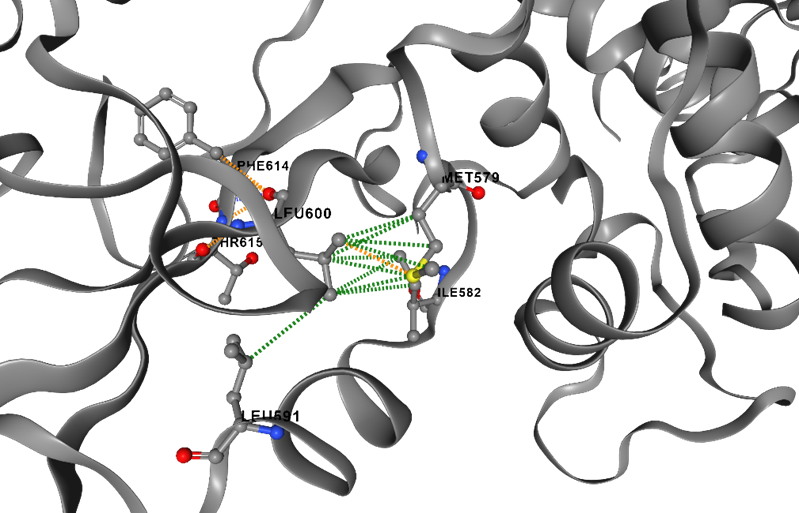

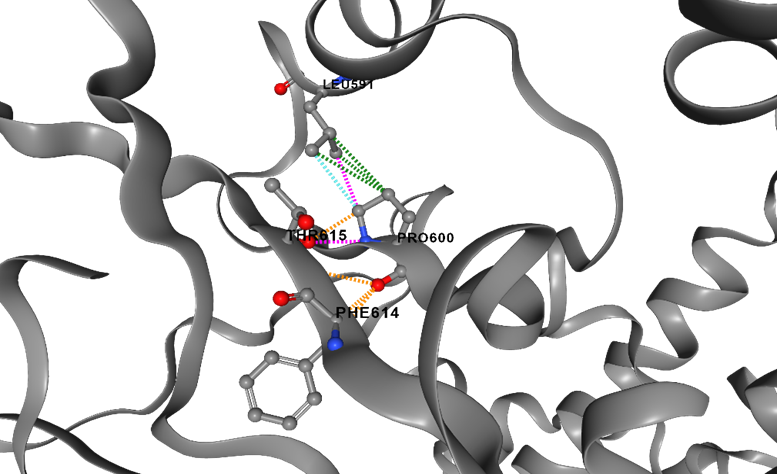

(A)L600P - wild type	(B) L600P mutant type

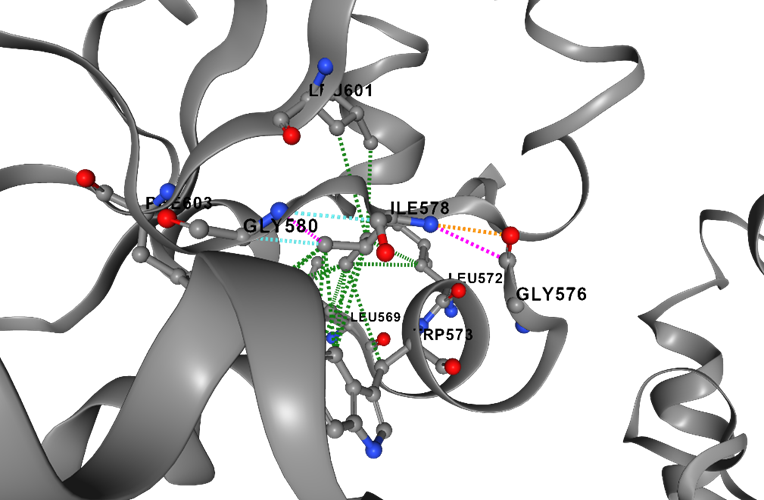

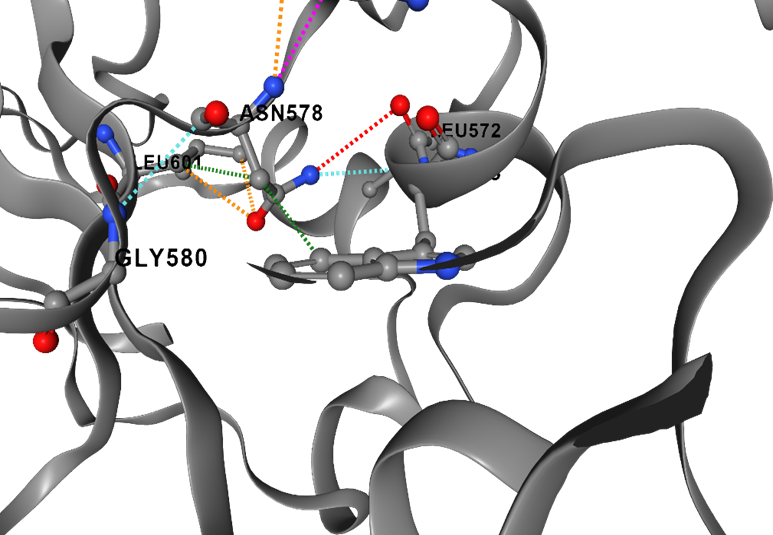

(A)I578N - wild type	(B) I578N, mutant residue

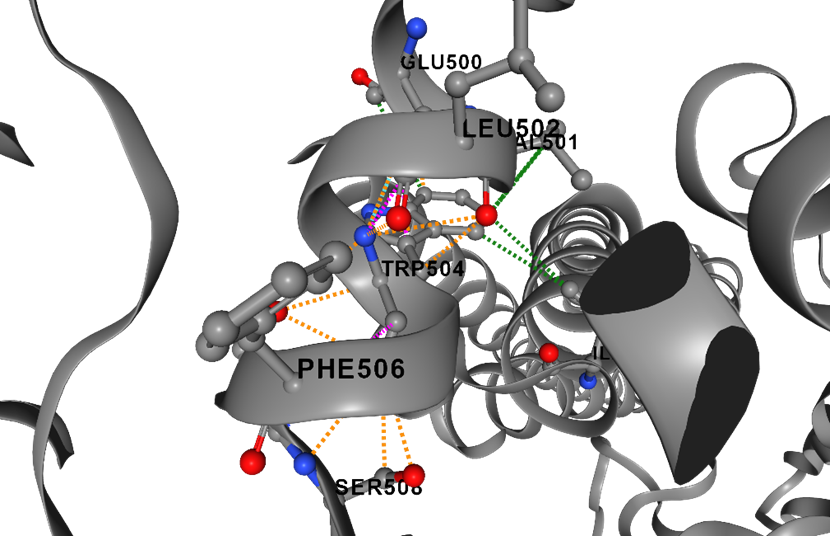

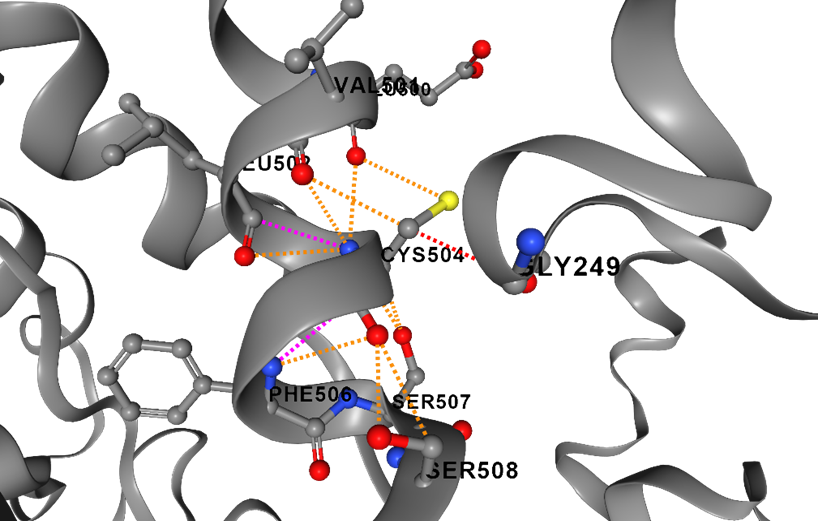

(A)W504C - wild type	(B) W504C, mutant residue

| Hydrogen Bond | Hydrophobic | VDW | Clash | Ionic | Aromatic | Polar | Carbonyl |
| --- | --- | --- | --- | --- | --- | --- | --- |

Figure S6. Difference in ionic interactions between the wild-type (A) and mutant residues (B) in I648T.
