## supplementary table 1 for "Integrating Artificial Intelligence and Bioinformatics Methods to Identify Disruptive STAT1 Variants Impacting Protein Stability and Function"

Table S1. List of nsSNPs that were predicted to have pathological significance by PhD-SNP, SNPs & GO, P Mut, and PANTHER.

|  | **nsSNP** | **Amino acid**  **change** | **P MUT** | | **PhD-SNP** | | **SNP&GO** | | **PANTHER** | |
| --- | --- | --- | --- | --- | --- | --- | --- | --- | --- | --- |
|  |  |  | **Prediction** | **Score** | **Prediction** | **RI** | **Prediction** | **RI** | **Effect** | **Preservation**  **Time** |
| 1 | rs1374373369 | D674V | Disease | 90% | Disease | 9 | Disease | 8 | probably damaging | 1036 |
| 2 | rs759271255 | I648T | Disease | 92% | Disease | 6 | Disease | 6 | probably damaging | 842 |
| 3 | rs752542806 | V642D | Disease | 87% | Disease | 5 | Disease | 8 | probably damaging | 455 |
| 4 | rs1209841496 | R602W | Disease | 93% | Disease | 8 | Disease | 3 | probably damaging | 1237 |
| 5 | rs137852678 | L600P | Disease | 93% | Disease | 9 | Disease | 8 | probably damaging | 1036 |
| 6 | rs767475430 | I578N | Disease | 93% | Disease | 8 | Disease | 6 | probably damaging | 1237 |
| 7 | rs916580554 | W504C | Disease | 91% | Disease | 6 | Disease | 8 | probably damaging | 750 |
| 8 | rs527393923 | T450M | Disease | 86% | Disease | 1 | Disease | 2 | probably damaging | 1036 |
| 9 | rs865962653 | S51L | Disease | 88% | Disease | 5 | Disease | 1 | probably damaging | 750 |
