## supplementary table 2 for "Integrating Artificial Intelligence and Bioinformatics Methods to Identify Disruptive STAT1 Variants Impacting Protein Stability and Function"

Table S2. MutPred probability values of deleterious and pathogenic nsSNPs identified in STAT1

|  | **nsSNP ID** | **Amino acid**  **change** | **MutPred 2 score** | **Affected PROSITE and ELM Motifs** | **Molecular mechanisms P-values <= 0.05** | **Probability** | **P-value** |
| --- | --- | --- | --- | --- | --- | --- | --- |
| 1 | rs1374373369 | D674V | 0.813 | - | Gain of Strand | 0 .26 | 0.04 |
|  |  |  |  |  | Gain of Acetylation at K673 | 0.25 | 0.01 |
| 2 | rs759271255 I648T | | 0.893 |  | Altered Stability | 0.16 | 0.02 |
| 3 | rs752542806 V642D | | 0.867. | ELME000063 ELME000085  ELME000147  ELME000155  ELME000220  ELME000233 | Altered Ordered interface | 0.35 | 4.2e-03 |
|  |  |  |  |  | Gain of Relative solvent  Accessibility | 0.30 | 7.3e-03 |
|  |  |  |  |  | Altered Transmembrane protein | 0.18 | 8.6e-03 |
|  |  |  |  |  | Altered DNA binding | 0.15 | 0.04 |
| 4 | rs1209841496 | R602W | 0.896 | -  ELME000328  ELME000052  ELME000062 | Gain of Strand | 0.27 | 0.02 |
|  |  |  |  |  | Altered Stability | 0.09 | 0.05 |
| 5 | rs137852678 | L600P | 0.965 | ELME000052  ELME000328 | Gain of Intrinsic disorder | 0.31 | 0.04 |
|  |  |  |  |  | Altered Stability | 0.28 | 6.6e-03 |
| 6 | rs767475430 | I578N 0.936 | | PS00008 | - | - | - |
| 7 | rs916580554 | W504C 0.807 | | ELME000197 | - | - | - |
| 8 | rs527393923 | T450M 0.373 | | - | - | - | - |
| 9 | rs865962653 | S51L | 0.665 | ELME000063ELME000147  ELME000336 | Altered transmembrane protein | 0.23 | 2.4e-03 |
