## supplementary table 3 for "Integrating Artificial Intelligence and Bioinformatics Methods to Identify Disruptive STAT1 Variants Impacting Protein Stability and Function"

Table S3. Deleterious and pathogenic ns SNPs were predicted to have significant decrease on protein stability by I-MUTANT 3.0 algorithm, MUpro, and DDMUT

|  | **SNP ID** | **Amino acid**  **change** | **I mutant 3** | | | **MUpro DDMUT** | |
| --- | --- | --- | --- | --- | --- | --- | --- |
|  |  |  | **Stability** | **RI** | **DDG**  **(kcal/mol)** | **Stability** | **DDG Stability DDG**  **(kcal/mol) (kcal/mol)** |
| 1 | rs759271255 | I648T | Decrease | 9 | -2.43 | Decrease | -2.4802937 Destabilizing -2.93 |
| 2 | rs752542806 | V642D | Decrease | 8 | -1.85 | Decrease | -1.8071037 Destabilizing -1.11 |
| 3 | rs1209841496 | R602W | Decrease | 3 | -0.20 | Decrease | -1.0486884 Destabilizing: -0.19 |
| 4 | rs137852678 | L600P | Decrease | 3 | -1.54 | Decrease | -1.6074419 Destabilizing -3.06 |
| 5 | rs767475430 | I578N | Decrease | 5 | -1.92 | Decrease | -0.98144877 Destabilising -0.84 |
| 6 | rs916580554 | W504C | Decrease | 8 | -1.41 | Decrease | -0.86533645 Destabilizing -0.73 |
