## supplementary table 4 for "Integrating Artificial Intelligence and Bioinformatics Methods to Identify Disruptive STAT1 Variants Impacting Protein Stability and Function"

Table S4. Shows Alpha-missense prediction of the pathogenic nsSNPs in STAT1

|  | **SNP ID** | **Substitution** | **Alpha-missense pathogenicity** | **Alpha-missense prediction** |
| --- | --- | --- | --- | --- |
| 1 | rs759271255 | I648T | 0.9875 | Likely Pathogenic |
| 2 | rs752542806 | V642D | 0.9916 | Likely Pathogenic |
| 3 | rs1209841496 | R602W | 0.9982 | Likely Pathogenic |
| 4 | rs137852678 | L600P | 0.9998 | Likely Pathogenic |
| 5 | rs767475430 | I578N | 0.9986 | Likely Pathogenic |
| 6 | rs916580554 | W504C | 0.9815 | Likely Pathogenic |
