## supplementary table 5 for "Integrating Artificial Intelligence and Bioinformatics Methods to Identify Disruptive STAT1 Variants Impacting Protein Stability and Function"

Table S5. Conservation profile of most damaging nsSNPs of STAT1

| **No** | **SNP ID** | **Amino acid**  **change** | **Conservation**  **score** | **Prediction** |
| --- | --- | --- | --- | --- |
| 1 | rs759271255 | I648T | 8 | Conserved and buried |
| 2 | rs752542806 | V642D | 6 | Buried |
| 3 | rs120984149 | R602W | 9 | (functional residues), highly conserved and exposed |
| 4 | rs137852678 | L600P | 9 | (structural residues), highly conserved and buried |
| 5 | rs767475430 | I578N | 9 | (structural residues), highly conserved and buried |
| 6 | rs916580554 | W504C | 8 | Conserved and buried |
