## supplementary table 6 for "Integrating Artificial Intelligence and Bioinformatics Methods to Identify Disruptive STAT1 Variants Impacting Protein Stability and Function"

Table S6. Changes in physical properties between wild-type and mutant residues predicted by project hope

|  | **SNPs** | **Differ in size** | **Difference in charge** | **Difference in hydrophobicity** | **Disrupt hydrogen bond** | **Affect contact with ligand molecules** |
| --- | --- | --- | --- | --- | --- | --- |
| 1 | I648T | Yes | No | Yes | No | Yes |
| 2 | V642D | Yes | No | Yes | No | Yes |
| 3 | R602W | Yes | No | Yes | No | Yes |
| 4 | L600P | Yes | No | Yes | No | No |
| 5 | I578N | Yes | No | Yes | Yes | Yes |
| 6 | W504C | Yes | No | Yes | No | Yes |
