## supplementary table 7 for "Integrating Artificial Intelligence and Bioinformatics Methods to Identify Disruptive STAT1 Variants Impacting Protein Stability and Function"

Table S7. Domain regions of the selected most damaging nsSNPs in STAT1

| **STAT1 domains (position)** | **SNPs** |
| --- | --- |
| STAT1, SH2 domain (557–707) | Y668F, I648T , V642D, R602W, and L600P |
| STAT1 transcription factor, DNA binding domain (323–458) | R304C |
| SH2 domain (578–638) | I578N |
| Src homology 2 (SH2) domain profile (573-670) | I578N |
